# Mechanistic Basis for Enhanced Strigolactone Sensitivity in KAI2 Triple Mutant

**DOI:** 10.1101/2023.01.18.524622

**Authors:** Briana L. Sobecks, Jiming Chen, Tanner J. Dean, Diwakar Shukla

## Abstract

*Striga hermonthica* is a parasitic weed that destroys billions of dollars’ worth of staple crops every year. Its rapid proliferation stems from an enhanced ability to metabolize strigolactones (SLs), plant hormones that direct root branching and shoot growth. *Striga’s* SL receptor, *Sh*HTL7, bears more similarity to the staple crop karrikin receptor KAI2 than to SL receptor D14, though KAI2 variants in plants like *Arabidopsis thaliana* show minimal SL sensitivity. Recently, studies have indicated that a small number of point mutations to HTL7 residues can confer SL sensitivity to *At* KAI2. Here, we analyze both wild-type *At* KAI2 and SL-sensitive mutant Var64 through all-atom, long-timescale molecular dynamics simulations to determine the effects of these mutations on receptor function at a molecular level. We demonstrate that the mutations stabilize SL binding by about 2 kcal/mol. They also result in a doubling of the average pocket volume, and eliminate the dependence of binding on certain pocket conformational arrangements. While the probability of certain non-binding SL-receptor interactions increases in the mutant compared with the wild-type, the rate of binding also increases by a factor of ten. All these changes account for the increased SL sensitivity in mutant KAI2, and suggest mechanisms for increasing functionality of host crop SL receptors.

## Introduction

Witchweed, also known as *Striga*, is a parasitic plant that destroys an estimated $10 billion of crops every year, which impacts around 100 million farmers worldwide.^1^ *Striga* proliferates by metabolizing strigolactones (SL), a hormone exuded from the roots of staple food crops, and germinates at nM or pM concentrations via activity of the *Sh*HTL protein family.^2–4^ The most sensitive of these is *Sh*HTL7, which germinates at SL concentrations as low as 2 pM. ^3,5^ The analogous receptor in host crops, KAI2, is only sensitive at micromolar concentrations, so understanding how high sensitivity evolved in *Sh*HTL7 may provide a means for targeting the receptor and inhibiting the destructive proliferation of *Striga*.^3^

KAI2, or karrikin insensitive 2, induces germination in plants such as *Arabidopsis thaliana* by metabolizing smoke-derived compounds called karrikins that contain a butenolide structure fused to a six-membered ring.^6–8^ SLs are a class of plant hormones linked to regulation of lateral root and branch development.^9–13^ SL molecules have two main moieties connected by an enol-ether bridge: a tricyclic portion (A, B and C rings) and the butenolide ring (D-ring).^12,14^ *Sh*HTL7, *At* KAI2, and *At* D14 (the SL receptor for *Arabidopsis*) are all α-β hydrolases with a conserved Ser-Asp-His catalytic triad. ^15–17^ However, while both karrikins and SLs contain a butenolide ring, *At* KAI2 is not sensitive to SLs.^6^ Phylogenetic analysis indicates that KAI2 paralogs fall into three different clades: KAI2c, the conserved form of the gene which includes *At* KAI2; KAI2i, the intermediate form of the gene; and KAI2d, the divergent form which includes *Sh*HTL7. KAI2d genes convergently evolved the same functionality as *At* D14, which split off from the KAI2 clade at an earlier date.^18^ A recent study found even more evidence supporting the emergence of SL sensitive KAI2d proteins from SL insensitive KAI2 proteins.^19,20^ Only 3 amino acid substitutions from WT *At* KAI2 to corresponding residues on *Sh*HTL7 (Trp153Leu, Phe157Thr, and Gly190Thr) induced SL sensitivity.^21^ Enhanced SL sensitivity in KAI2d versus other proteins correlates with an observed increase in pocket volume and flexibility in *Sh*HTL7 relative to other KAI2-related proteins.^3,21^ The specific mutant here, known as Var64, also showed a 10-fold increase in time spent in the productive binding mode with a corresponding decrease in the unproductive binding mode.^21^ There was also an increase in downstream signaling partner gene expression in strains of *Arabidopsis* containing Var64.^21^ However, the pocket volume and binding calculations were performed on either static structures or small-scale molecular dynamics (MD) simulations (on the order of 0.5-1 µs). These studies do not give a complete view of receptor dynamics or functionality across long timescales. For long-timescale processes like ligand binding and receptor activation, a much greater quantity of simulation data is needed to accurately characterize the thermodynamics and kinetics of the system. ^22,23^

MD simulation is a powerful technique for characterizing long-timescale processes of proteins,^25^ such as ligand binding of the SL molecule to receptor proteins, with full-atom resolution.^26^ Employing Markov State Models (MSMs) in combination with MD simulation limits sampling bias and offers accurate kinetic and thermodynamic characterization of a system.^27^ MSMs are a method for assigning millions of frames in an MD system to a small number of kinetically relevant conformations and computing the rates of interconversion between them. Previously, we have performed long-timescale MD simulations with MSMs to characterize plant hormone perception including strigolactone and associated conformational change processes.^26,28–35^ In this study, we created two systems to study the difference in molecular interactions between the WT and Var64 and how these differences provide a mechanism for SL metabolism in the latter but not the former.^21^ Specifically, we sought to see the effect of the three mutations on SL-residue interactions, pocket volume, and binding kinetics. We ran one system with the wild-type version of *At* KAI2 and one with the Var64 mutant (Trp153Leu, Phe157Thr, and Gly190Thr), with about 200 µs of aggregate simulation data for each system. Both systems contained one molecule of GR24, a synthetic SL analogue.^36^ Initial simulations were performed using adaptive sampling, in which we clustered simulation frames based on structural features, selected undersampled regions based on a “least-counts” principle, and generated subsequent simulations using frames from the undersampled regions as a starting point. The adaptive sampling and least counts approach efficiently characterizes a system by directing future simulations towards areas with less data and avoiding areas that have already been adequately sampled.^37–40^ Once enough data was generated to cover a sufficient portion of the landscape, the rest of the samples were run on the Folding@Home distributed computing system (for further details, see Methods section in SI).^41,42^ In our data analysis, we investigated both thermodynamic effects of the mutations to examine any stabilization of poses conducive to binding and kinetic effects to see any increase of flux into ligand-bound states. In this paper, we show that the triple mutant of *At* KAI2 improves thermodynamic and kinetic properties. These mutations promote more stable SL binding, enhance contacts that stabilize a wider pocket volume, increase anchoring of GR24 near the pocket entrance, and contribute towards a tenfold increase in the binding flux.

## Methods

### Molecular Dynamics Simulations

#### Simulation Protocol

Wild-type (WT) and mutant systems were created from structure 4IH1 from the Protein Data Bank.^24^ Amino acid mutations were added to the mutant system using Tleap in Amber18.^43^ ACE and NME terminal caps were added using CHARMM-GUI (Residues 1 and 270 for D14).^44–46^ The GR24 model was taken from a bound structure of *Os*D14 (PDB 5DJ5) and was inserted using Packmol into a random position away from the receptor in both the WT and mutant systems.^47,48^ Simulations were set up with AmberTools 18 and run with Amber 18 using the ff14SB force field.^43,49^ Water was described with the TIP3P model and GR24 with the generalized AMBER force field (GAFF).^50,51^ The simulation box size was a 70 Å cube containing TIP3P water and an NaCl concentration of 0.15 M to provide a neutral charge.^52^ Structures were minimized with the conjugate gradient descent method for 10000 steps, then equilibrated for 5 ns. The Langevin thermostat kept simulations at a constant temperature of 300 K and the Monte-Carlo barostat kept a constant pressure of 1.0 bar. Short-range non-bonded interactions were calculated with a cutoff of 10 Å and long-range electrostatics calculated with the Particle Mesh Ewald algorithm.^53^ The SHAKE algorithm was used for constraining bonds to hydrogen.^54^ After three rounds of simulations (2.6 µs) in Amber, 1500 independent runs for both the WT and mutant systems were then performed using OpenMM on the Folding@Home distributed computing system starting from the conformations obtained by clustering the initial simulation data into 100 states. ^41,55^

#### Adaptive Sampling

To generate enough points for the Folding@Home simulations, we employed an adaptive sampling scheme known as least-counts. Following an initial round of simulations, we clustered all frames around a few known structural metrics, determined the least sampled conformations, and used these conformations to seed the next rounds. After enough data was generated to sample the landscape, extensive simulations for both the wild-type and mutant systems were run using the Folding@Home distributed computing system. ^41^ In total, 26.65 µs of data was generated for the adaptive sampling rounds and 200 µs of data was generated for both the wild-type and mutant system from Folding@Home. Sampling metrics and a summary of simulations run for each round are provided in Tables S1 and S2.

### Trajectory Analysis

#### Feature Calculations

All contact information was calculated with the MDTraj analysis package version 1.9.4.^56^ All residue contact distances were calculated with residue alpha carbons except for contacts between mutated residues and residue F26, which were computed using the closest heavy sidechain atom. GR24 features were calculated with individual carbons in either the A-ring or the D-ring.

#### Pocket Volume Calculation

Pocket volume was calculated with Epock version 1.0.5.^57^ Volume was calculated within a “maximum englobing region” (MER) defined as a sphere centered at the midpoint between the S95 Cα and the geometric center of the α carbons of the T1-T2 helices (resid 138 to 160). The MER radius was defined as the distance between the sphere center and the S95 Cα. Measured pocket volumes were weighted with MSM probabilities.

#### Contact Probability Calculation

Contact probability was calculated for GR24 against all residues in the WT and mutant KAI2. Contact probability for a given residue is defined as the fraction of total frames in which GR24 is within 4 Å of the residue. Contact probabilities were weighted by MSM probabilities (*π_i_*) using Eq. 1.

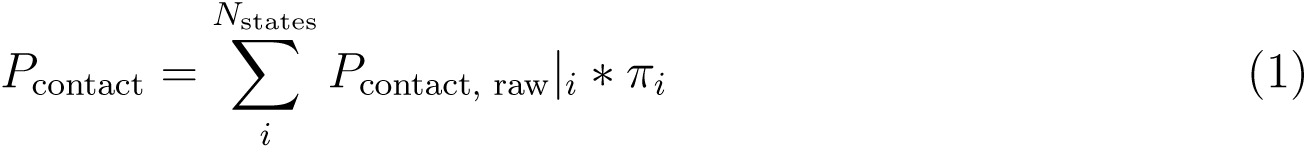

Overall probabilities of ligand binding were calculated using

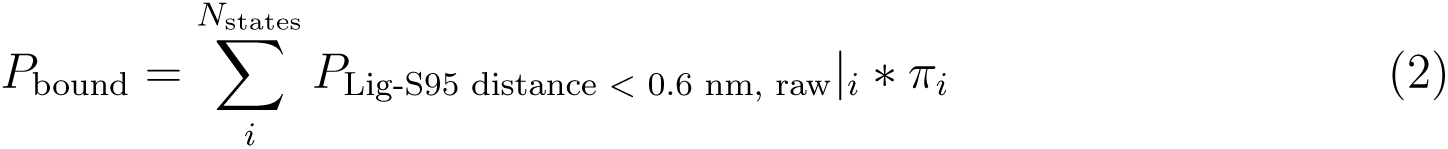

Probability of ligand binding was also used to represent the probability of the ligand being present in the pocket. The probability of the ligand at the pocket entrance followed a similar formula, but used the ten residues on the T1 and T2 helix that face inward towards the pocket in the KAI2 crystal structure (residues 139, 142, 143, 146, 147, 153, 154, 157, 158, and 161).

### Markov State Model Construction

Markov state models were built in pyEMMA from 41 total metrics related to ligand binding, mutant residue interactions, and catalytic triad activity.^58,59^ A complete list of metrics is available in the Supporting Information (Table S3). A hyperparameter search was performed using cross-validation to generate a VAMP1 score.^60^ Final MSM hyperparameters are listed in Table S4. Markovian behavior of MSMs was evaluated using the Chapman-Kolmogorov test (Fig. S3 and S4). Using the state equilibrium probabilities from Markov state models, free energy landscapes were calculated using

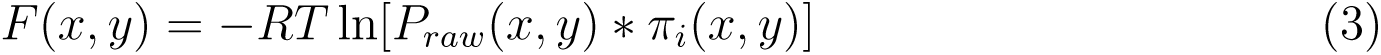

### Transition Path Theory

Transition path theory is a method to calculate the flux between a small number of kinetically relevant macrostates as a means of estimating which states have the highest rate of interconversion.^61,62^ Four macrostates for the system were defined based on the position of GR24 with respect to KAI2. These states were bound, unproductively bound, anchored, and unbound. State definitions are given in Table S5. The mean first passage time (MFPT) was computed between each pair of macrostates. The MFPT gives the amount of time required to reach state B for the first time when starting at state A. Transition path theory and the MFPT highlight which states are accessed quickest and allow a clear comparison between the kinetic profile of two systems. ^26^

## Results and Discussion

### Var64 shows enhanced binding to SLs compared with WT KAI2

To explore the experimental finding of enhanced SL binding in Var 64 versus the wild type, we investigated several different properties of WT *At* KAI2 and the Var64 mutant by calculating different structural metrics of the systems.^21^ Figure 2 shows the ligand binding behavior of both the WT and Var64. In KAI2 homologs, catalysis is believed to occur when the S95 residue nucleophilically attacks the SL molecule’s D-ring. ^63,64^ Figures 2(A) and 2(B) show the free energy landscape projected onto A-ring and D-ring distances from S95 in the WT and Var64 systems, respectively. Both systems show high stability in the (α) position, corresponding to the bound state of KAI2 (D-ring facing the pocket). However, Var64 shows a higher population of bound states, and the stability of states can reach up to 2-3 kcal/mol lower than in the wild-type (Fig. 2(B)). We calculated the probability of binding, or the probability that GR24 is within a certain distance of the catalytic serine (*<* 0.6 nm for binding), and found this probability to be 0.017 in the WT and 0.038 in the mutant. Therefore, binding is enhanced more than twofold in mutant versus wild-type *At* KAI2. Both states also show a high degree of stability in an “inverse bound” state, where the A-ring instead of the D-ring is oriented towards the pocket (β). However, there is a lower free energy barrier between the bound and inverse states in Var64 versus the wild-type. Additionally, the free energy of the unbound state denoted by δ in Figure 2A-B is approximately 2 kcal/mol higher for Var64, showing that SL binding is more favorable in the mutant protein. This binding is also affected by the residues involved in the mutation. In the WT, the probability of GR24 being in contact with W153 is 0.159 and with F157 is 0.167, but these probabilities increase in Var64 to 0.168 for L153 and 0.203 for T157, respectively (see Fig. 4). When examining a visual representation of SL binding generated from these contact probabilities, a degree of steric hindrance is observed between GR24 and the bulky aromatic wild-type residues W153 and F157 (Fig. 2(C)). In contrast, the mutant residues L153 and T157 are smaller and inhibit less of the binding pocket volume (Fig. 2(D)).

**Figure 1:**
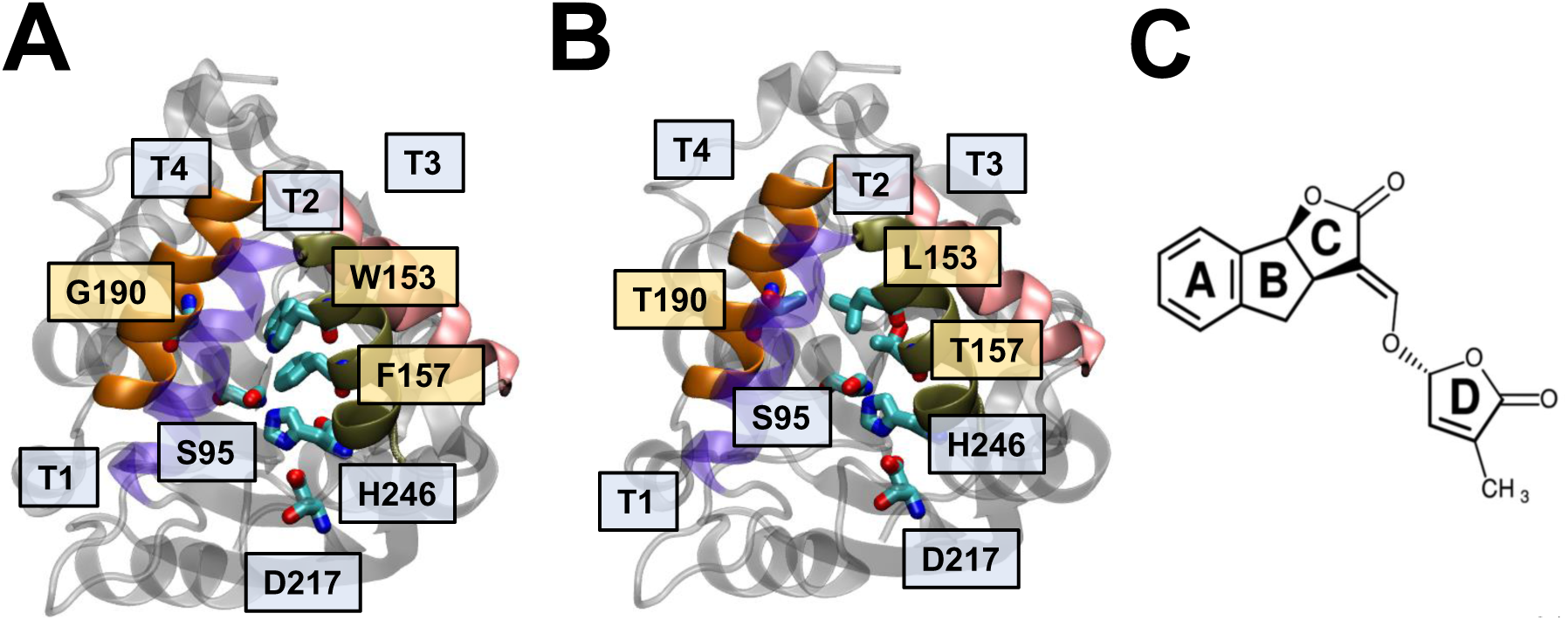
Key structural features of (A) wild-type (WT) *At* KAI2 (PDB code 4IH1^24^), (B) Var64 *At* KAI2 mutant, (C) and strigolactone analogue GR24. Major helices on both *At* KAI2 variants are T1, shown in indigo, T2, shown in tan, T3, shown in pink, and T4, shown in orange. Important residues are depicted in cyan. S97, H247, and D218 make up the catalytic triad, which binds GR24. The three residues of interest for the mutation study are W153, F157, and G190 in the WT protein (A), and the mutated versions are L153, T157, and T190 (B). The mutant residue labels are in yellow boxes, while other features are in blue boxes. (C) The structure of GR24, labeled with appropriate ring structures. This is the R-form of GR24, which is the chirality exhibited by naturally occurring strigolactones.^12^

**Figure 2:**
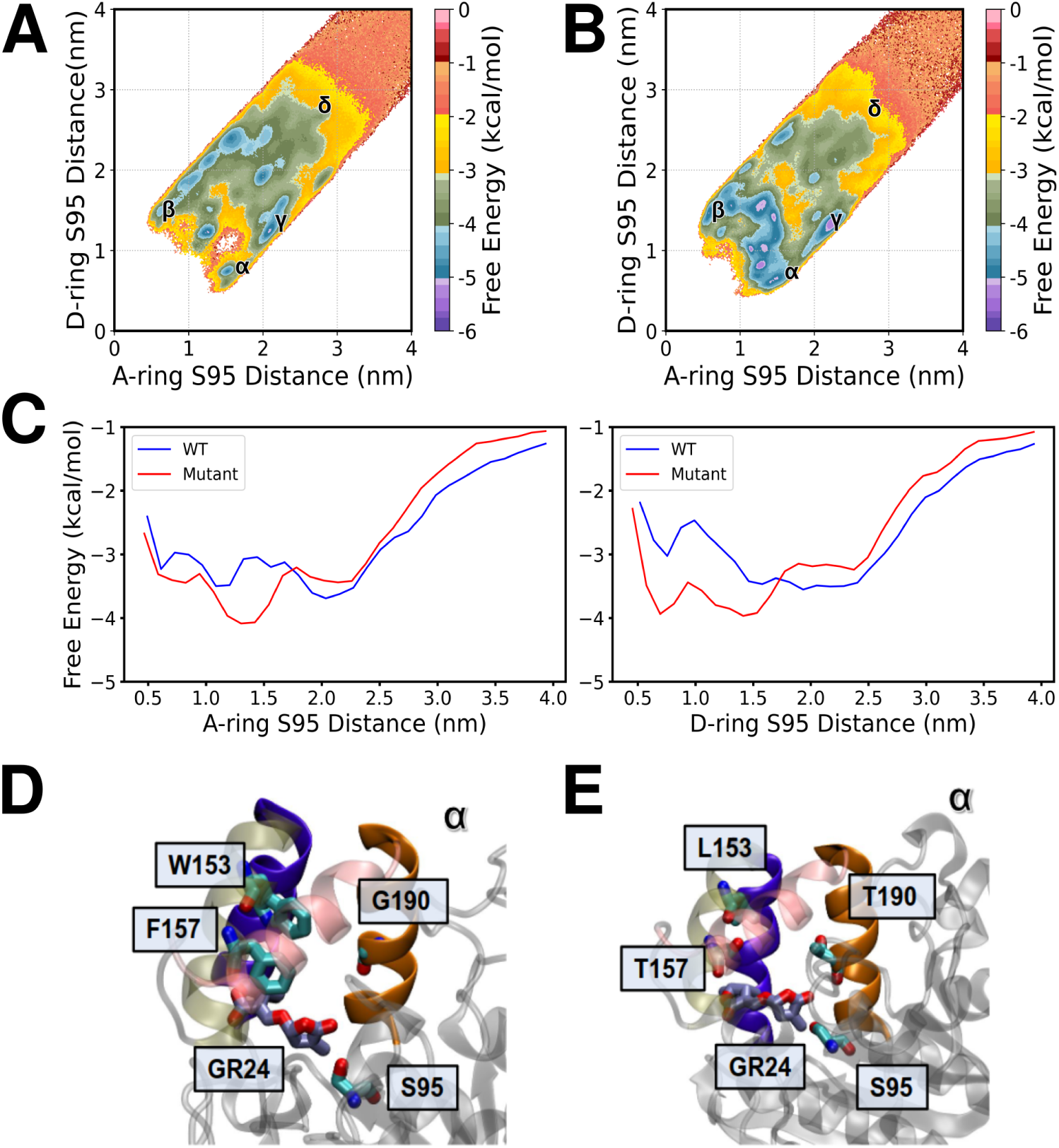
SL binding in WT and mutant *At* KAI2. MSM-reweighted binding free energy landscapes for WT (A) and mutant (B) systems. α denotes the productively bound state (D-ring facing S95), β the unproductively bound state (A-ring facing S95), γ the lid-helix anchored state, and δ the unbound states. (See SI Fig. S5 for error plots). (C) 1D free energy projections of the A-ring S95 and D-ring S95 distances. (D) GR24 binding in WT and (E) mutant systems.

### Var64 demonstrates a higher pocket volume versus WT

Given the observed difference in pocket free space in the representative binding images, we sought to characterize the pocket volume across both systems with a more rigorous quantitative approach. Previous work has shown a correlation between increased pocket volume and increased sensitivity to SLs, as well as a positive correlation between pocket flexibility and SL sensitivity.^21^ We calculated the average pocket volume for the WT protein as 357 Å^3^ and for the mutant protein as 678 Å^3^, which is nearly double that of the WT. Figure 3(A) shows a close approximation of the average pocket volume of the wild-type protein, and Figure 3(B) shows a close approximation of the average volume of the mutant. After computing the true average, we selected all simulation frames within a 1 Å^3^ range of that average and chose one at random. We also found the probability distribution of the WT and mutant pocket volumes (Figure 3(C)). The two proteins showed a similar distribution around their means, with a standard deviation of 134 Å^3^ for the WT and 146 Å^3^ for the mutant. Therefore, the flexibility of the mutant is only marginally higher than the wild-type, but the center of the distribution is higher in Var64. The change in T2 helix residues 153 and 157 from bulky aromatics to smaller side-chain amino acids is therefore likely a major reason for the difference in pocket volume. The significantly greater binding pocket volume in Var64 suggests a strong reason for the enhanced binding in mutant *At* KAI2, with enhanced pocket flexibility playing a smaller but still visible role.

**Figure 3:**
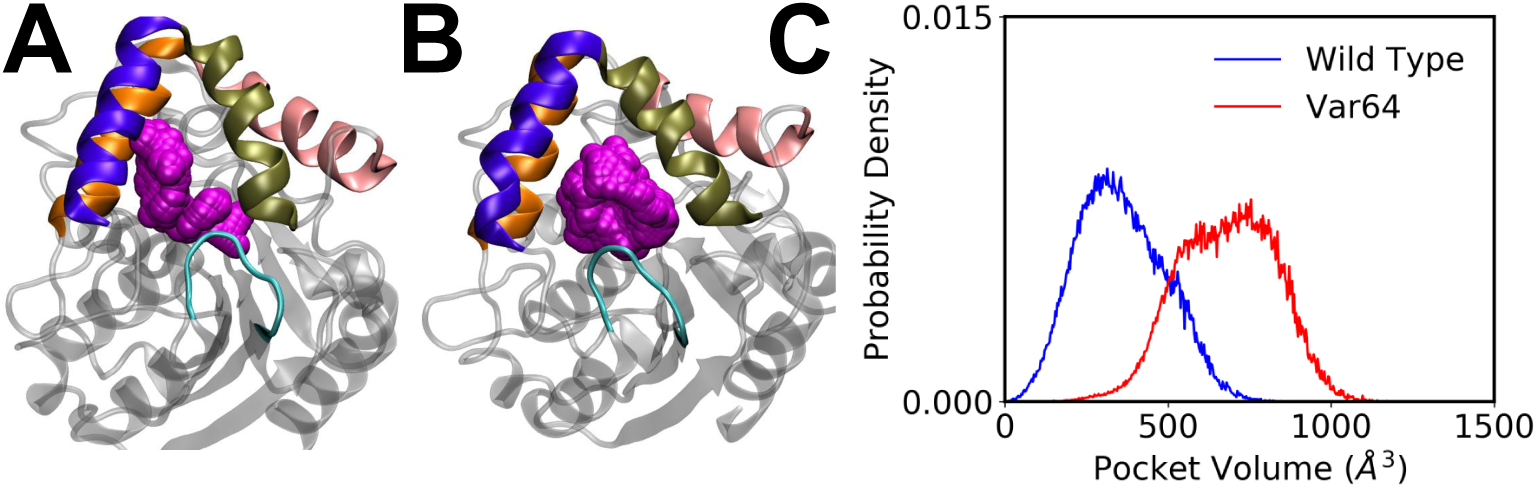
Volume of the binding pocket in WT and Mutant *At* KAI2. (A) Approximate av-erage volume of the WT pocket. Free space is given in magenta spheres. The structure was chosen by calculating the average pocket volume, then selecting all conformationally generated structures within 1 Å^3^ of the average and selecting one at random to be a representative structure. (B) Average volume of the mutant pocket chosen using the same procedure as in (A). (C) Probability distribution of WT and Var64 mutant pocket volume.

**Figure 4:**
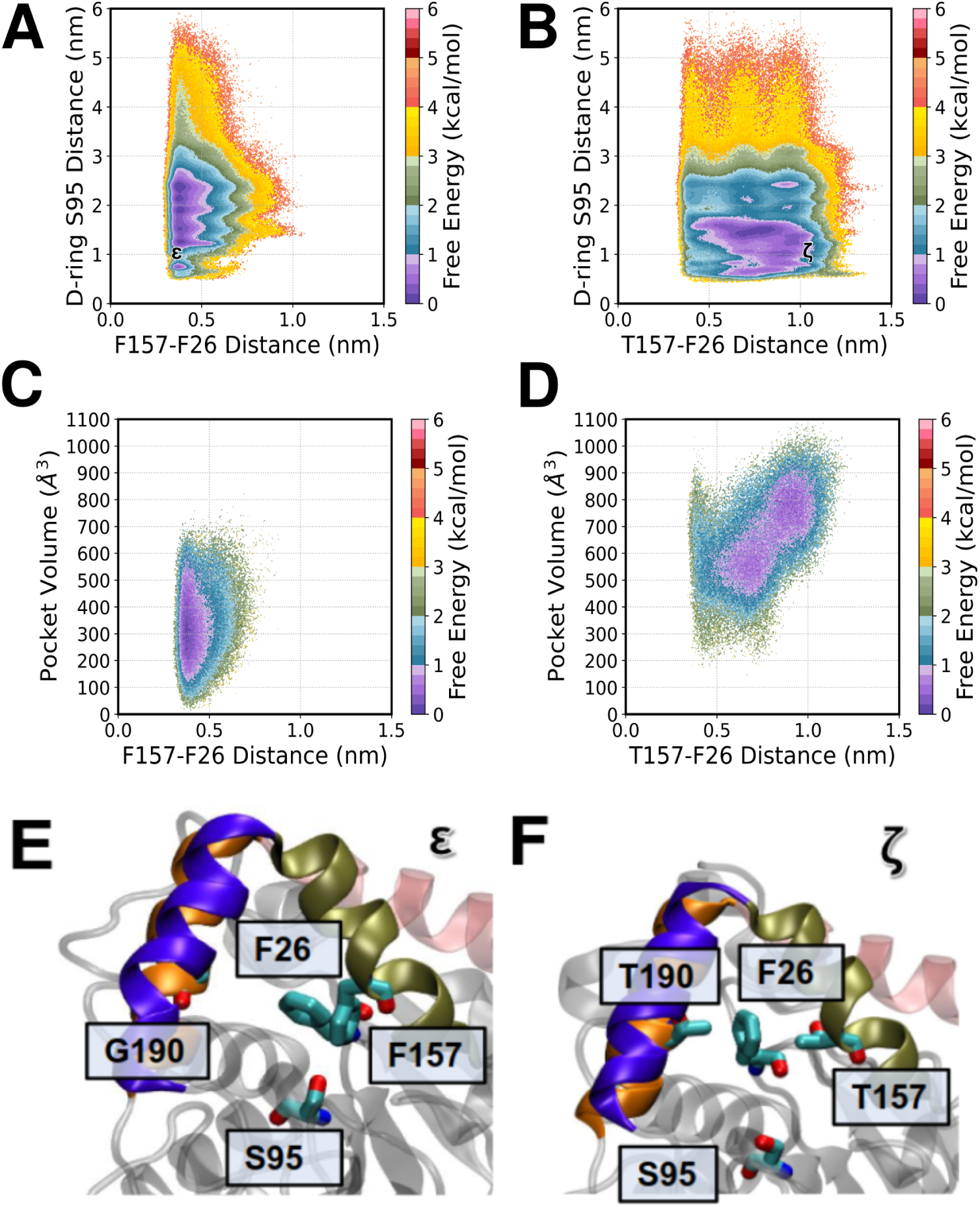
Pocket dynamics of WT and mutant *At* KAI2. (A) Free energy landscape of WT F157-F26 interaction (T2 helix to back of pocket) versus SL binding. (B) The same interaction but with the mutant T157 residue. ε denotes a point of high T2 helix-back of pocket interaction. (C) Free energy landscape of WT F157-F26 interaction versus pocket volume. (D) The same interaction but with the mutant T157 residue. ζ indicates a point of high T4 helix-back of pocket interaction. (See SI Fig. S6 for error plots). (E) Close F157-F26 interaction in WT *At* KAI2 (ε). (F) Separated T157-F26 residues in mutant *At* KAI2 (ζ).

### Var64 and WT lid helix contact differences affect binding stability

After characterizing the pocket volume of both systems, we investigated any contacts responsible for affecting the volume of the WT versus mutant system. One residue of note was F26, found on a loop at the back of the binding pocket. In preliminary data prior to MSM-weighting, F26 was in contact with GR24 for about a quarter of all simulation frames, leading to further investigation of the residue’s behavior (Figure S7). In the wild-type system, the lowest free energy wells in the system were located where the distance between the closest sidechain F26 and WT residue F157 was less than 5 Å (Fig. 4(A)). The system expresses the highest stability when these two residues are in close contact with each other. In contrast, the mutant residue T157 displayed a wider range of distances from F26 (Fig. 4(B)). While the D-ring only gets close enough for binding (*<* ∼0.6 nm according to the binding cutoff from our previous work) when F157 and F26 are in close contact, binding is independent of the distance between T157 and F26 in the mutant.^26^ We also examined the relationship between F26 and residue 157 distance versus pocket volume. In the WT system, there is no correlation between F157-F26 distance on pocket volume (Fig. 4(C)), but there is a clear linear relationship between these two variables in the mutant (Fig. 4(D)). Given that S95 binding is independent of T157-F26 distance, ligand binding can occur when the two residues are far apart and the pocket is wider. This gives a more accessible volume for GR24 to enter the pocket and bind, in agreement with the findings in previous experimental work with the mutant.^21^ All of these interactions serve to demonstrate the difference in pocket behavior between WT and Var64 *At* KAI2. Since the T3 helix spreads out from the other helices (Fig. 4(E) and 4(F)), the T2 helix is open for direct contact with the back of the pocket. The high degree of contact between F26 and F157, from hydrophobic contact between the two residues that draws the loop at the back of the pocket towards the T2 helix, leaving less volume in the pocket and blocking space for the WT to interact with catalytic S95 (Fig. 4(E)). When F26 is further from F157, the loop stays further back in the pocket, giving more space for binding (Fig. 4(F)). Additionally, these interactions keep F26 in an upright position with its aromatic ring vertical. Even when interacting with T157, F26 keeps this orientation, versus in the WT, where it bends forward (Fig. 4(E)). All of these interactions display a demonstrable increase in the mutant pocket’s volume and provide more space for SL binding.

### Contact probability of GR24 with WT and mutant

The difference in pocket conformations between WT and mutant *At* KAI2 led us to investigate the overall ligand contacts with the entire protein in both systems through contact probability analysis. Contact probability is defined as the fraction of frames in which the ligand is within a certain cutoff distance, in this case 4 Å, of a given residue. This distance is the approximate cutoff for the length scale of van der Waals interactions. ^26^ We computed the MSM-reweighted contact probability for every residue across our entire simulation trajectories for both systems. Figure 5(A) shows the contact probability of GR24 with all WT residues. The points of highest contact were around the T1 and T2 helices (residues 138-160). Of the 270 residues in KAI2, the one with the highest contact was F157, occurring in 16.7 ± 0.01% of frames. Figure 5(B) gives a representation of WT KAI2 with contact probabilities mapped onto the structure, highlighting the high degree of contact between GR24 and the residues of interest in the mutagenesis study. Figure 5(C) displays the contact probability of the Var64 mutant. While a large portion of this graph resembles the wild-type one, the contact probabilities of T1 and T2 helix residues is significantly higher. Here, the GR24 contact probability with residue T157 is 20.3 ± 0.02% of all frames. The increase in contact probabilities here likely results from the increase in time spent near the binding pocket, though GR24 is not in contact with these residues while in the bound state (Figure S6). The projection of probabilities onto the mutant structure is given in Figure 5(D). We also computed a “pocket entrance probability”, or the probability that GR24 was in contact with any of the ten amino acids facing the pocket interior in the KAI2 crystal structure (see Methods section for a complete list). The WT pocket entrance probability was 32.1 ± 0.02% and the mutant value was 43.2 ± 0.03%, following the trend of increased anchoring observed in Var64 versus wild-type KAI2. Finally, we directly compared the difference in contact probability between the two systems (Figure 5(E)). The difference between Var64 and the WT in the lid helix region was around 5 ± 0.02% for many residues. These difference values are projected onto Figure 5(F). As described previously, the probability of binding was more than two times greater in the mutant than the wild-type protein. Therefore, while the “anchored” state probability is one-third times higher in the mutant protein, the bound state probability experiences a greater proportional increase in the mutant.

**Figure 5:**
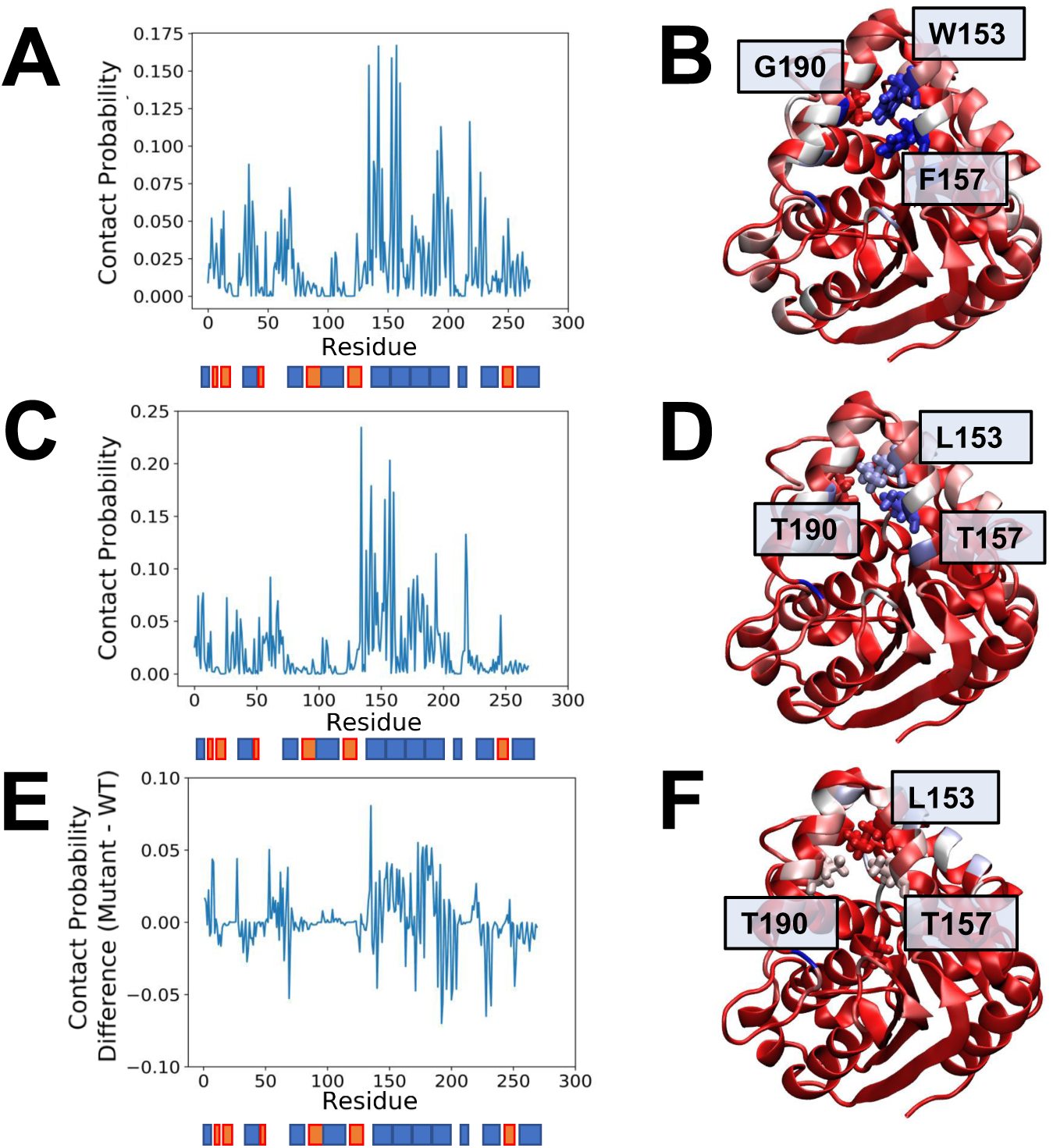
Contact probability of GR24 against WT and mutant residues. Contact probability is the fraction of frames in which the ligand interacts with a given residue. Contact probability is given for the wild-type (A), mutant (C), and the difference between the two (E). A secondary structure map is given below the graph, with alpha helices in blue, and beta sheets in orange. (B) Contact probability of WT residues mapped onto the KAI2 crystal structure. High-contact residues are blue, medium-contact are white, and low-contact are red. Coloring for the degree of contact is relative for each individual structure. The 3 residues changed in the mutant are shown in licorice representation. (D) Contact probability of mutant residues mapped onto the mutated KAI2 crystal structure, with the same representation as (B). (F) Contact probability difference mapped onto the mutated KAI2 crystal structure. Blue and white indicates a greater degree of GR24 contact with mutant *At* KAI2, while red indicates no difference or greater contact with the WT.

### Transition path theory

We performed a transition path theory analysis to see the relationship between various states of GR24 binding activity. Figure 6 shows the relative fluxes between states in the WT (A) and mutant (B) systems. The rate from unbound and unproductively bound states to the bound state was about ten times higher in the mutant than in the WT (on the order of 10 vs 1 µs). The rate from the anchored state to the bound state was about thirteen times higher in the mutant. Therefore, it seems that it is more likely in the mutant for the ligand to both find the binding pocket from free solution and from other interaction positions with KAI2. The rate of GR24 dissociation from productive binding, moreover, was nearly twice as high in the WT as in the mutant, indicating that GR24 is less likely to unbind when bound to the mutant. However, GR24 in the mutant was twice as likely to go from the productively bound to unproductively bound position compared with the WT, and over three times more likely to go from the bound state to the anchored state. Given the higher binding pocket volume of Var64, the ligand could have more flexibility to move easily between states in the mutant. Both of these processes were on the order of 100 microseconds, much higher than the rate of binding. Still, the large relative increase in binding flux for the mutant corroborates the increased SL sensitivity of Var64.

**Figure 6:**
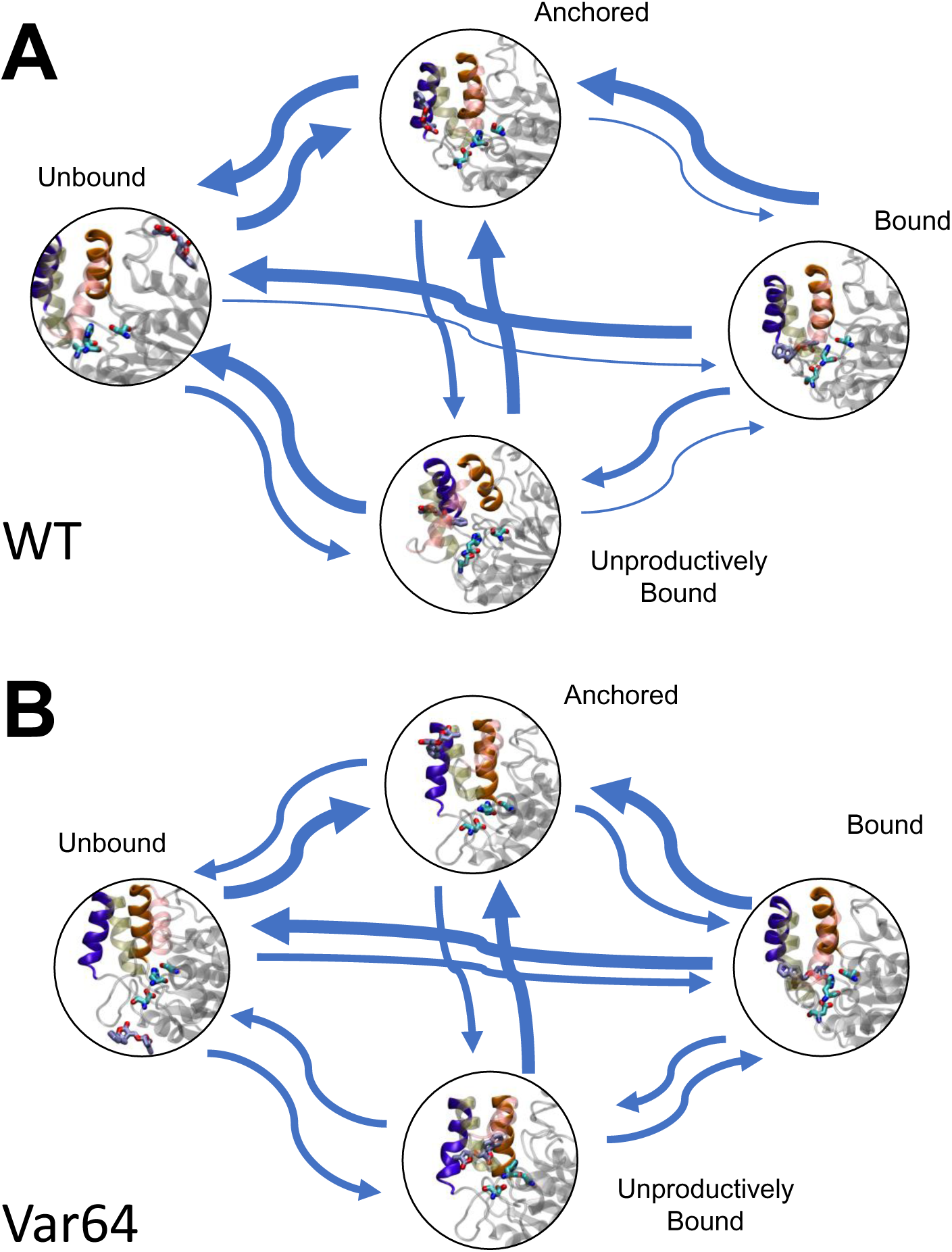
Transition path theory of WT (A) and mutant (B) binding. Arrow thickness represents the relative flux between two states. The four states defined here are productive binding, where the D-ring of GR24 is oriented towards the catalytic residues; unproductive binding, where the A-ring is oriented towards the catalytic residues; anchoring, where GR24 associates with the T1 and T2 helices; and unbound, where the ligand is not in any other relevant position. Thin arrows represent processes on the scale of 1 µs, medium arrows represent 10 µs, and thick arrows 100 µs. Exact values can be found in Table S6.

## Conclusions

At the molecular level, the three amino acid substitutions in the Var64 *At* KAI2 mutant have a profound impact on SL sensitivity. GR24 binding was approximately 2 kcal/mol more stable in the mutant versus the WT, and the binding region was more populated as well. When examining the pocket behavior, we found that the average pocket volume in Var64 was nearly double that of the WT, and the pocket was marginally more flexible, indicating that more space is available for entering the pocket and binding to the catalytic residues. Part of this could be explained by the replacement of bulky aromatic sidechains with smaller ones, but that alone does not account for the difference. An investigation of mutant residue contacts within the binding pocket found that the WT version of residues needed to occupy a precise arrangement for binding to occur, but the residue contacts had much greater flexibility in the mutant, especially in the contact between T157 and F26. While allowing for greater pocket flexibility, however, mutations also promoted an increase in lid helix anchoring of GR24. However, while the flux to lid-anchored states from bound states tripled in the mutant, the flux from lid-anchored to bound states increased by a factor of thirteen, showing that this lid anchoring enhanced binding more than it detracted from it. The flux to bound from unbound states increased ten times in the mutant as well, showing that the mutations indeed have a profound effect on binding kinetics. Overall, these three mutations enhance SL binding at the molecular level, suggesting a probable evolutionary path for SL perception in *Striga*. Our findings provide a key for understanding the mechanisms involved in SL perception and explaining the heightened sensitivity in proteins such as *Sh*HTL7 relative to host crop analogues.

## Supporting Information

This article contains supporting information. PDF document containing additional information describing methods and validation, including adaptive sampling details, Markov state model construction parameters and validation, and transition path theory definitions and raw data; video of ligand binding pathway.

## Data Availability

All full-atom trajectories for both the WT and KAI2 mutant, and analysis scripts can be found at the following Box link: https://uofi.box.com/s/zvchi4sqx09ql662olwv6hrftdw5tp3a.

## Author Contributions

D.S. acquired funding for the project. D.S. and J.C. conceptualized and supervised the project. B.L.S. and J.C. performed simulations and analyzed data. B.L.S. and J.C. wrote the original manuscript draft with inputs from D.S. T.J.D performed analysis, data curation, and revised the manuscript in response to the reviewer comments. All authors reviewed and edited the manuscript.

## Acknowledgement

B.L.S. acknowledges support from the XSEDE EMPOWER program under National Science Foundation grant number ACI-1548562 and the Clare Boothe Luce scholars program. J.C. is a member of the NIH Chemistry-Biology Interface Training Program (T32-GM136629).

D.S. acknowledges support from the CAS Fellowship, Center for Advanced Studies at University of Illinois at Urbana-Champaign and Sloan Research Fellowship from Alfred P. Sloan Foundation. All three acknowledge support from the Folding@Home distributed computing system and its users for donating computing power necessary for running simulations for this projects.

