## Supplementary Information for "Mechanistic Basis for Enhanced Strigolactone Sensitivity in KAI2 Triple Mutant"

### Details of adaptive sampling protocol

Table S1: Adaptive sampling metrics and selection criteria for both systems

| Round | Sampling Metrics | Selection Criteria |
|---|---|---|
| 1 | - | Initial structure |
| 2 | S95-GR24 A-ring distance<br>S95-GR24 D-ring distance | 4 random clusters, 25 points per cluster<br>and D-ring-S95 distance $< 5$ nm |
| 3 | S95-GR24 A-ring distance<br>S95-GR24 D-ring distance | 100 clusters,<br>5 points each from 20 least populated<br>and D-ring-S95 distance $< 5$ nm |
| FAH | S95-GR24 A-ring distance<br>S95-GR24 D-ring distance | 100 clusters,<br>5 points each from 20 least populated<br>and D-ring-S95 distance $< 3$ nm |

Table S2: Summary of adaptive rounds for both systems

| Round | Parallel Trajectories | Trajectory length (ns) | Aggregate ($\mu$s) |
|---|---|---|---|
| 1 | 5 | 130 | .650 |
| 2 | 100 | 130 | 13.1 |
| 3 | 100 | 130 | 12.9 |
| | | Total ($\mu$s) | 26.65 |

### Markov state model construction and validation

#### MSM Features

To construct our Markov state models, we first calculated a set of inter-residue distance and helical content features from our simulation data. The full set of features used for MSM construction is shown in Table S3.

Table S3: Featurizations used for MSM construction. C-$\alpha$ distances were used for all inter-residue distances.

| | WT | Mutant |
|---|---|---|
| Catalytic Triad | S95-D217 | S95-D217 |
| | S95-H246 | S95-H246 |
| | D217-H246 | D217-H246 |
| D-Loop-H246-Distances | V215-H246 | V215-H246 |
| | K216-H246 | K216-H246 |
| | L218-H246 | L218-H246 |
| | A219-H246 | A219-H246 |
| | V220-H246 | V220-H246 |
| | P221-H246 | P221-H246 |
| A-ring-D-Loop-Distances | D217-A ring | D217-A ring |
| | A219-A ring | A219-A ring |
| | P221-A ring | P221-A ring |
| D-ring-D-Loop-Distances | D217-D ring | D217-D ring |
| | A219-D ring | A219-D ring |
| | P221-D ring | P221-D ring |
| Ligand - Catalytic Triad | A-ring-S95 | A-ring-S95 |
| | D-ring-S95 | D-ring-S95 |
| Ligand - Mut Resids | D-ring-W153 | D-ring-L153 |
| | D-ring-F157 | D-ring-T157 |
| | D-ring-G190 | D-ring-T190 |
| Mut Resids | W153-F157 | L153-T157 |
| | W153-G190 | L153-T190 |
| | F157-G190 | T157-T190 |
| T1-T2 Distances | D139-L160 | D138-L160 |
| | L142-F157 | L142-T157 |
| | I146-W153 | I146-L153 |
| A-ring T1 Distances | A-ring-K139 | A-ring-K139 |
| | A-ring-L142 | A-ring-L142 |
| | A-ring-A145 | A-ring-A145 |
| A-ring T2 Distances | A-ring-W153 | A-ring-L153 |
| | A-ring-F157 | A-ring-T157 |
| | A-ring-L160 | A-ring-T160 |
| D-ring T1 Distances | D-ring-K139 | D-ring-K139 |
| | D-ring-L142 | D-ring-L142 |
| | D-ring-A145 | D-ring-A145 |
| D-ring T2 Distances | D-ring-W153 | D-ring-L153 |
| | D-ring-F157 | D-ring-T157 |
| | D-ring-L160 | D-ring-L160 |

#### MSM hyperparameter selection

To select lag times for our MSMs, we plotted implied timescales for models estimated at a series of lag times and chose a lag time at which the implied timescale stopped changing with increasing lag time. Implied timescale plots are shown in Fig. S1.

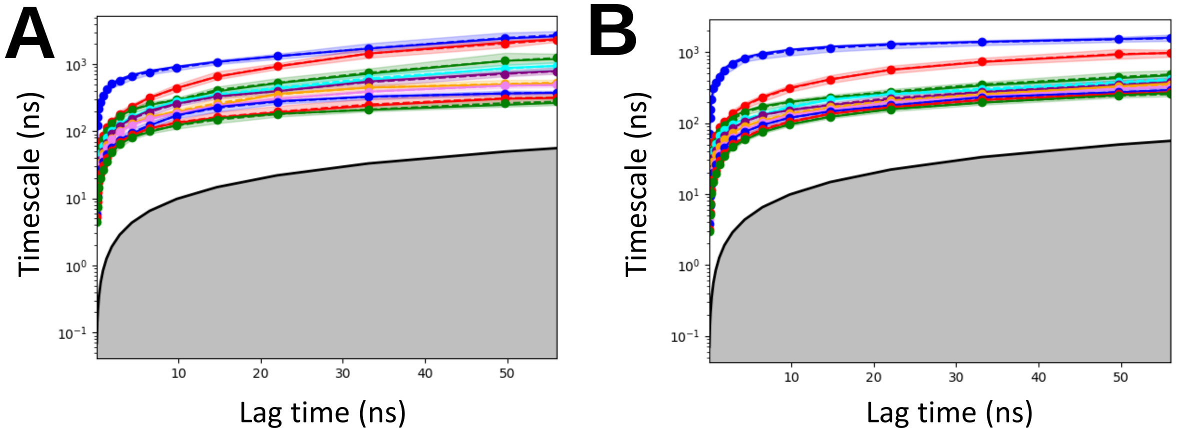


Figure S1: Implied timescale plots for the WT (A) and mutant (B) systems. Lag time was chosen as 15 ns for both systems.

To select number of TICA components and number of clusters to discretize our simulation data for MSM construction, we performed a grid search in which we calculated a cross-validation score for MSMs calculated with different parameter sets. Cross-validation scores for different parameter sets are shown in Fig. S2. Final parameters used for MSM construction are shown in Table S4.

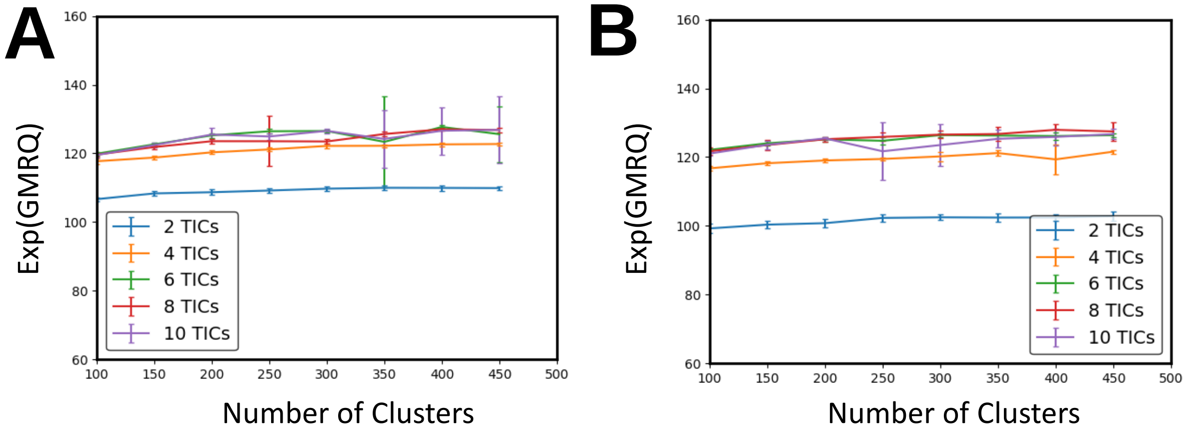


Figure S2: Cross-validation scores calculated with different hyperparameter sets for the WT and mutant (B) systems.

Table S4: Final parameters used for MSM construction

| | WT | Mutant |
|---|---|---|
| Lag time (ns) | 15 | 15 |
| Number of TICA components | 6 | 6 |
| Number of clusters | 300 | 150 |

#### Chapman-Kolmogorov validation

To validate our MSMs, we employed the Chapman-Kolmogorov test. Briefly, if a system exhibits Markovian behavior, the $n$th power of a transition matrix estimated at lag time $\tau$ should equal the transition matrix estimated at lag time $n\tau$. Results of the Chapman-Kolmogorov test are shown in Fig. S4.

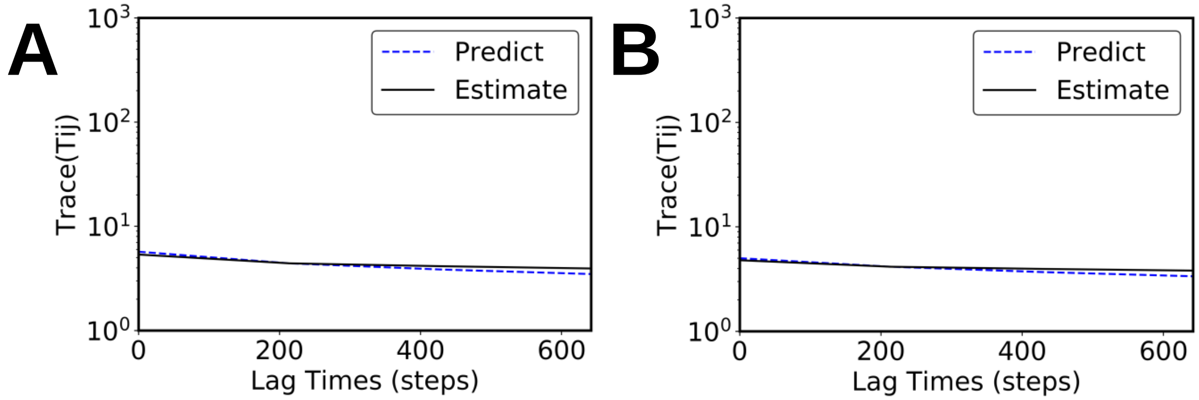


Figure S3: Chapman-Kolmogorov tests for the WT (A) and mutant (B) systems. The test shows good agreement between estimated and predicted MSMs, indicating that the models follow the Markov property.

We also performed the Chapman-Kolmogorov test on five metastable states. Here too, we see a good degree of agreement between the estimated and predicted coarse-grained transition matricies.

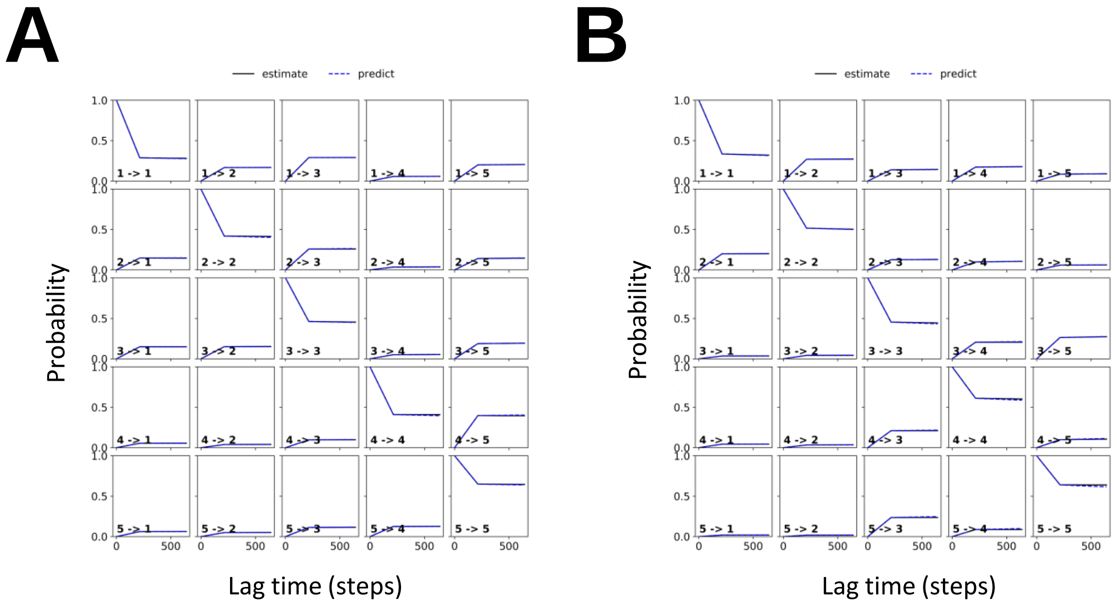


Figure S4: Chapman-Kolmogorov tests of five metastable states for the WT (A) and mutant (B) systems, demonstrating Markovian behavior.

#### SL binding free energy landscape error

We computed free energy errors using a bootstrapping procedure where we built an MSM using 80% of our data, calculated the mean and standard deviation, and repeated the process five times.

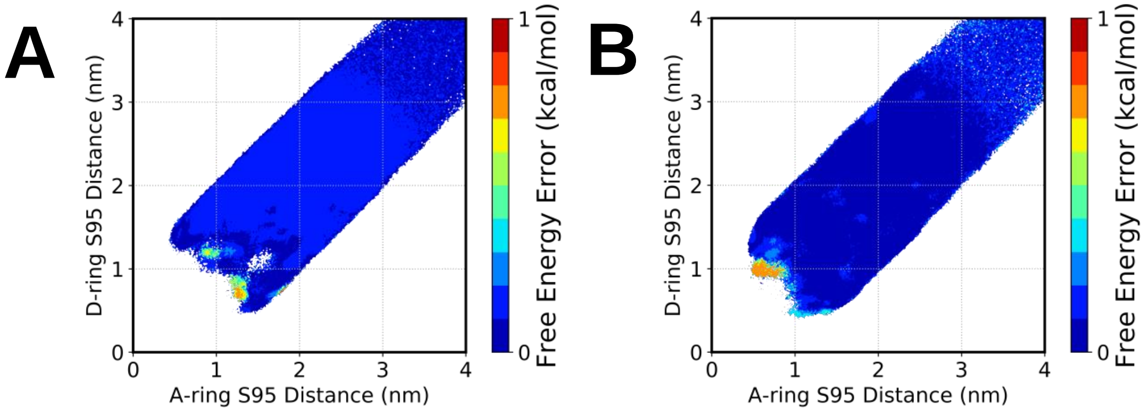


Figure S5: Free energy landscape error of SL binding for the WT (A) and mutant (B) systems.

#### SL F26 Contacts

As in the previous section, we computed free energy errors using a bootstrapping procedure where we built an MSM using 80% of our data, calculated the mean and standard deviation, and repeated the process five times.

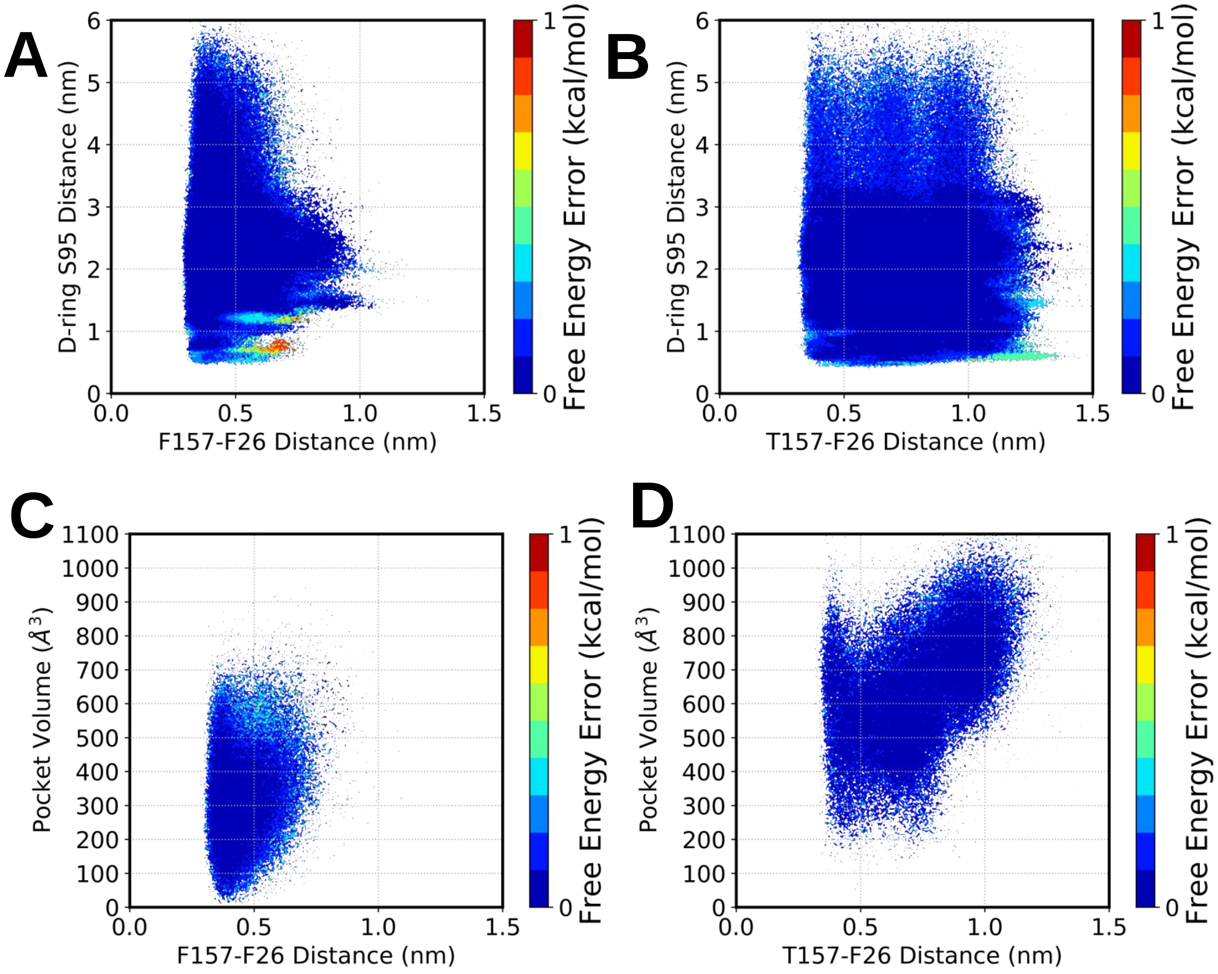


Figure S6: Free energy landscape error of F26 contacts versus GR24 D-ring binding for the WT (A) and mutant (B) systems and versus pocket volume for the WT (C) and mutant (D) systems.

### Preliminary contact probability

Before MSM weighting our systems, we examined preliminary contact probabilities and found a region of high contact with residue F26. The preliminary unweighted data is shown here.

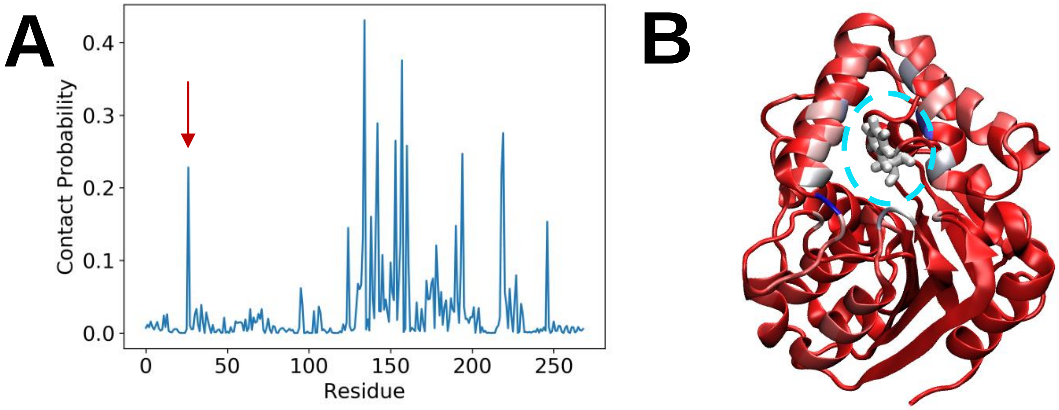


Figure S7: (A) Unweighted contact probability for the mutant system. The red arrow designates residue F26. (B) Unweighted contact probability projected onto the mutant crystal structure. F26 is shown in licorice representation and is surrounded by a light blue dashed circle.

### Binding versus anchoring

To confirm there was no overlap between ligand contact with the lid helices in the anchored state and ligand contact with the catalytic serine, we plotted a free energy landscape comparing the position of GR24 relative to these two features in the WT and mutant. These landscapes demonstrate that ligand contact with the lid helices and with the catalytic serine are mutually exclusive.

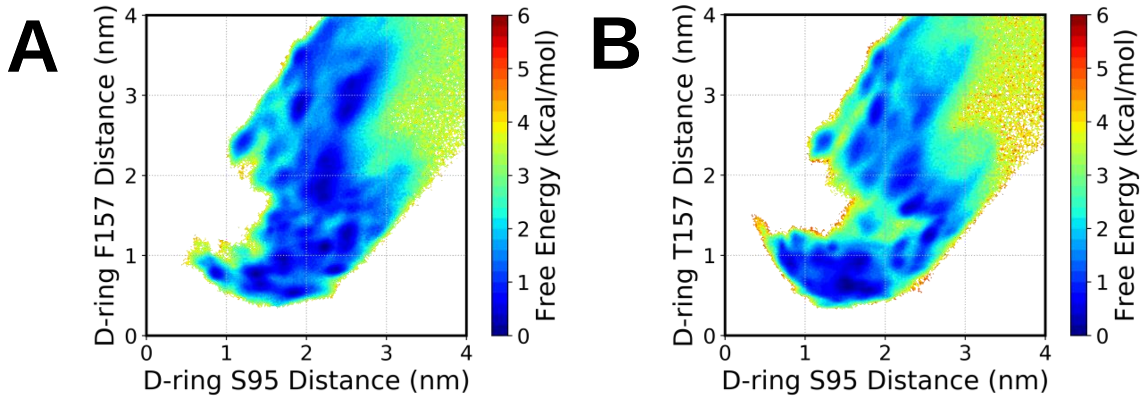


Figure S8: Free energy landscapes for ligand binding versus lid helix anchoring in the WT (A) and mutant (B) systems. These landscapes show that GR24 does not contact the catalytic serine and the lid helices at the same time.

### Transition path theory

During transition path theory calculation for the *holo*, we coarse-grained our data by assigning MSM clusters to a set of four macrostates defined by the position of the GR24 ligand. Definitions for these positions are shown in Table S5. Calculated relative flux between these four macrostates are shown in Table S6.

Table S5: Definitions of ligand binding and CTH association used for TPT calculation

| State | Parameters |
|---|---|
| Productive bound | Minimum D-ring-S95 distance $<$ 0.6 nm<br>AND<br>Mean A-ring-S95 distance $>$ Mean D-ring-S97 distance |
| Unproductive bound | Mean A-ring-S95 distance $<$ 1.5 nm<br>AND<br>Mean D-ring-S95 distance $<$ 1.5 nm<br>AND<br>Not included in anchored state |
| Anchored | Minimum A-ring-A145 distance $<$ 0.9 nm<br>AND<br>Minimum A-ring-W153/L153 distance $<$ 0.9 nm<br>AND<br>Minimum D-ring-K139 distance $<$ 0.9 nm<br>AND<br>Minimum D-ring-L160 distance $<$ 0.9 nm<br>AND<br>Not included in bound state |
| Unbound | Minimum A-ring-S95 distance $<$ 1.5 nm<br>AND<br>Minimum D-ring-S95 distance $<$ 1.5 nm<br>AND<br>Not included in any other state |

Table S6: Relative flux values in microseconds (μs) as calculated by transition path theory

| | WT | Mutant |
|---|---|---|
| Bound → Anchored | 244.41 | 894.91 |
| Bound → Unbound | 188.64 | 110.06 |
| Bound → Unproductive bound | 35.49 | 67.82 |
| Anchored → Bound | 6.76 | 91.05 |
| Anchored → Unbound | 269.75 | 50.98 |
| Anchored → Unproductive bound | 88.66 | 87.83 |
| Unbound → Bound | 6.53 | 56.64 |
| Unbound → Anchored | 237.69 | 188.62 |
| Unbound → Unproductive bound | 47.22 | 58.73 |
| Unproductive bound → Bound | 5.69 | 56.77 |
| Unproductive bound → Anchored | 347.51 | 374.00 |
| Unproductive bound → Unbound | 149.20 | 34.43 |
